# Structural basis of streptomycin off-target binding to human mitoribosome

**DOI:** 10.1101/2022.02.02.478878

**Authors:** Yuzuru Itoh, Anas Khawaja, Vivek Singh, Andreas Naschberger, Joanna Rorbach, Alexey Amunts

## Abstract

The ribosome in mitochondria regulates cellular energy production, and its deactivation is associated with pathologies and ageing. Inhibition of human mitoribosome can be caused by antimicrobial off-target binding, which leads to clinical appearances. The anti-tuberculosis drug aminoglycoside streptomycin targets the small subunit and was shown to be coupled with a bilateral decreased visual acuity with central scotomas and an altered mitochondrial structure. Previously, we reported mitochondria-specific aspects of translation related to specialties of the human mitoribosome (Aibara et al., 2020). In this Research advance article, we report 2.23-Å resolution structure of the human mitoribosomal small subunit in complex with streptomycin. The structural data reveals new details of the streptomycin interactions, including specific water molecules and metal ions involved in the coordination. The density for the streptose moiety reveals that previously modeled aldehyde group appears as a loosely bound density, and the hydroxyl group is not resolved. The density replacing the aldehyde group is within hydrogen bonding distance of four phosphate groups of rRNA, suggesting that the ribosome-bound streptomycin is likely to be in the hydrated gem-diol form rather than in the free aldehyde form. Since streptomycin is a widely used drug for treatment, the newly resolved fine features can serve as determinants for targeting.

## Introduction

Human mitoribosome has a distinct structure, it receives mRNAs from Leucine-rich PPR-motif-containing protein (LRPPRC) and synthesizes 13 respiratory chain proteins delivered to the inner mitochondrial membrane via the OXA1L insertase (Aibara et al., 2020; Itoh et al., 2021). Dysfunction of human mitoribosome can be caused by off-target binding of antimicrobials that act on protein synthesis. This can lead to clinical symptoms of deafness, neuropathy, and myopathy, however the phenomenon can also be used to suppress glioblastoma stem cells growth (Sighel et al., 2021), and thus repurposing of mitoribosome-targeting antibiotics offers a therapeutic option for tumors (Vendramin et al., 2021). The aminoglycoside streptomycin that targets a ribosomal small subunit (SSU) was shown to be coupled with a bilateral decreased visual acuity with central scotomas and an altered mitochondrial structure (Kogachi et al., 2019). Moreover, patients carrying mtDNA mutations in the 12S rRNA gene, such as 1555A>G or 1494C>T are more prone to aminoglycoside-induced ototoxicity (Gao et al., 2017). To minimize toxic off-target effects, the approaches based on *in silico* modeling employing high resolution single-particle cryo-EM structures can be used. Although the sensitivity of mitoribosomes to antimicrobials has been documented, no detailed structural information elucidating specific molecular interactions is available, thus mechanistic details remained unknown.

## Results and discussion

To characterize the binding under close to physiological conditions, we added streptomycin to growing human embryonic kidney 293T (HEK293T) cells at a final concentration of 100 μg/ml and not to any of the biochemical purification steps. This approach implies that the antimicrobial would have to be imported into mitochondria, and therefore has an advantage over *in vitro* complex formation, as specific modifications and more native inhibitory properties would be preserved. The mitochondria were isolated, and the SSU was purified and subjected to a cryo-EM analysis. Monosome and LSU particles were removed during 2D classification, and the remaining particles underwent auto-refinement and 3D classification with local angular search with a solvent mask to remove poorly aligned particles, refinement and Contrast transfer function (CTF) refinement including beam-tilt, per-particle defocus, per-micrograph astigmatism in RELION 3.1 (Zivanov et al., 2020). Particles were then separated into multi-optics groups based on acquisition areas and date of data collection. Second round of CTF refinement (beam-tilt, trefoil and fourth-order aberrations, magnification anisotropy, per-particle defocus, per-micrograph astigmatism) was performed, followed by 3D auto-refinement. Finally, to improve the local resolution, local-masked 3D auto-refinements were systematically applied (Figure 1—figure supplement 1).

The resulting structure of the SSU shoulder with bound streptomycin was determined at 2.23 Å nominal resolution (Figure 1—figure supplement 1 and Table 1). This represents a substantial improvement of the X-ray crystal structures at 3.00-3.45 Å resolution of the *in vitro* formed complexes with *T. thermophilus* ribosome (Carter et al., 2000; Demirci et al., 2013). The high resolution allowed us to report a more precise binding mode of streptomycin, including coordinated water molecules, and a chemical alteration that could not be previously detected (Figure 1).

**Table 1:**
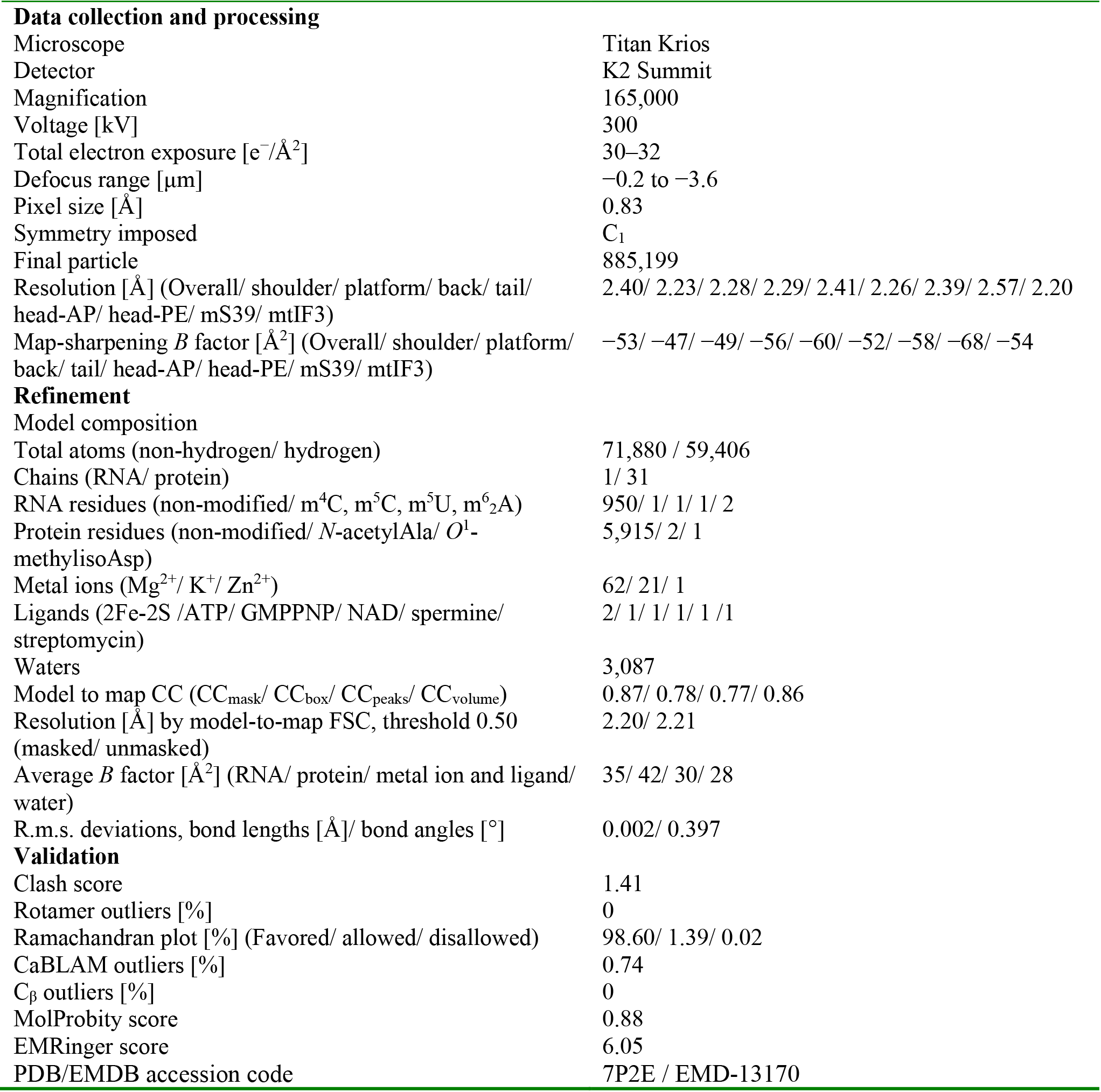
Cryo-EM collection, processing, model refinement and validation statistics.

|  |  |
| --- | --- |
| **Data collection and processing** |  |
| Microscope | Titan Krios |
| Detector | K2 Summit |
| Magnification | 165,000 |
| Voltage [kV] | 300 |
| Total electron exposure [$e^-/\text{\AA}^2$] | 30–32 |
| Defocus range [$\mu\text{m}$] | −0.2 to −3.6 |
| Pixel size [$\text{\AA}$] | 0.83 |
| Symmetry imposed | $C_1$ |
| Final particle | 885,199 |
| Resolution [$\text{\AA}$] (Overall/ shoulder/ platform/ back/ tail/ head-AP/ head-PE/ mS39/ mtIF3) | 2.40/ 2.23/ 2.28/ 2.29/ 2.41/ 2.26/ 2.39/ 2.57/ 2.20 |
| Map-sharpening $B$ factor [$\text{\AA}^2$] (Overall/ shoulder/ platform/ back/ tail/ head-AP/ head-PE/ mS39/ mtIF3) | −53/ −47/ −49/ −56/ −60/ −52/ −58/ −68/ −54 |
| **Refinement** |  |
| Model composition |  |
| Total atoms (non-hydrogen/ hydrogen) | 71,880 / 59,406 |
| Chains (RNA/ protein) | 1/ 31 |
| RNA residues (non-modified/ $m^4\text{C}$, $m^5\text{C}$, $m^5\text{U}$, $m^6_2\text{A}$) | 950/ 1/ 1/ 1/ 2 |
| Protein residues (non-modified/ $N$-acetylAla/ $O^1$-methylisoAsp) | 5,915/ 2/ 1 |
| Metal ions ($\text{Mg}^{2+}$/ $\text{K}^+$/ $\text{Zn}^{2+}$) | 62/ 21/ 1 |
| Ligands (2Fe-2S /ATP/ GMPPNP/ NAD/ spermine/ streptomycin) | 2/ 1/ 1/ 1/ 1 /1 |
| Waters | 3,087 |
| Model to map CC ($\text{CC}_{\text{mask}}$/ $\text{CC}_{\text{box}}$/ $\text{CC}_{\text{peaks}}$/ $\text{CC}_{\text{volume}}$) | 0.87/ 0.78/ 0.77/ 0.86 |
| Resolution [$\text{\AA}$] by model-to-map FSC, threshold 0.50 (masked/ unmasked) | 2.20/ 2.21 |
| Average $B$ factor [$\text{\AA}^2$] (RNA/ protein/ metal ion and ligand/ water) | 35/ 42/ 30/ 28 |
| R.m.s. deviations, bond lengths [$\text{\AA}$]/ bond angles [$^\circ$] | 0.002/ 0.397 |
| **Validation** |  |
| Clash score | 1.41 |
| Rotamer outliers [%] | 0 |
| Ramachandran plot [%] (Favored/ allowed/ disallowed) | 98.60/ 1.39/ 0.02 |
| CaBLAM outliers [%] | 0.74 |
| $C_\beta$ outliers [%] | 0 |
| MolProbity score | 0.88 |
| EMRinger score | 6.05 |
| PDB/EMDB accession code | 7P2E / EMD-13170 |

**Figure 1.**
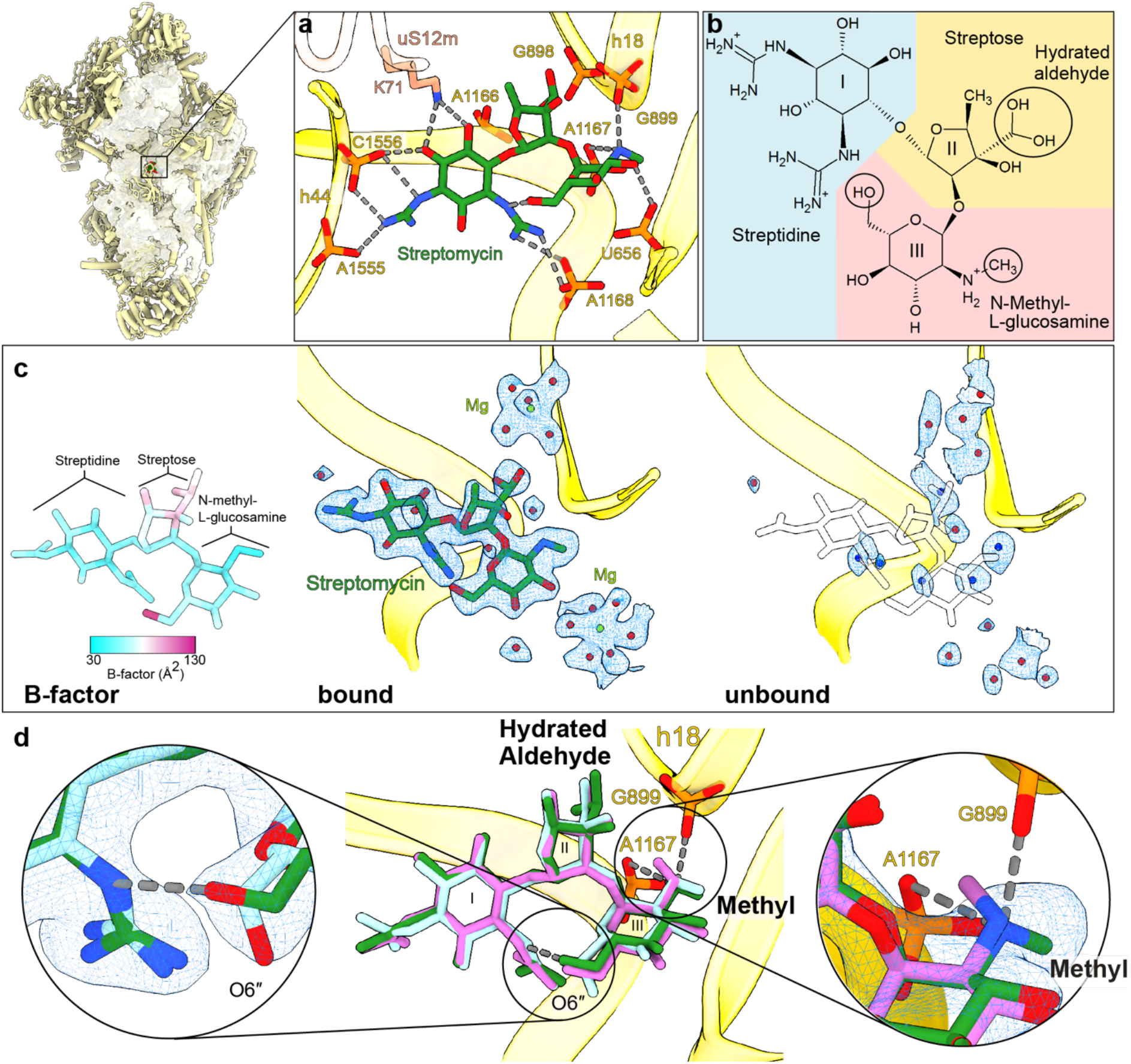
High resolution features of streptomycin binding to mitoribosome. (a) Streptomycin interacts with uS12m and backbone phosphates of helices h18 and h44. (b) Chemical structure of the hydrated gem-diol form of streptomycin. (c) Left, atomic *B*-factor distribution of streptomycin bound to our structure shows higher relative flexibility of the streptose moiety. Middle, density map and model of streptomycin along with surrounding water molecules and Mg^2+^ ions; Right, solvation of the binding site in the absence of streptomycin (indicated by a transparent frame). Water molecules present in absence or presence of streptomycin are colored red, while those observed only in the absence of streptomycin are colored blue. (d) Comparison of bound streptomycin in the current model with previously reported structures of bacterial ribosome (PDB ID: 4DR3, green; PDB ID: 1FJG, cyan) indicates that the model is supported by the density, and chemical interactions of O6′′ and methyl moieties resolve discrepancies in the previous models.

The chemical structure of streptomycin is comprised of three components linked by ether bonds: streptidine (scyllo-inositol with two hydroxyl groups substituted by guanidino groups), streptose (3-formyl-4-methyl tetrose), and *N*-methyl-*L*-glucosamine (Figure 1a,b). The density for the streptose moiety reveals a series of unexpected features (Figure 1c): 1) it is generally the poorest resolved component of streptomycin, 2) the methyl group of streptose is not well covered by the density, 3) the previously modeled aldehyde group appears as a loosely bound density, 4) the hydroxyl group is not resolved. Atomic *B*-factors estimated by reciprocal space refinement further support the idea that the streptose moiety is relatively flexible. The density replacing the aldehyde group is within hydrogen bonding distance of four phosphate groups of rRNA (C898, G899, A1166, A1167). Given that aldehyde has no hydrogen to provide for H-bonding phosphates, the ribosome-bound streptomycin is likely to be in the hydrated gem-diol form rather than in the free aldehyde form. The hydration is consistent with the NMR study of the free unbound state in aqueous solution (Blundell et al., 2013). Since the gem-diol group can H-bond with only two phosphate groups out of four, it suggests that the stabilization is not optimized, and this further explains the flexibility of the streptose moiety. Interestingly, in the biosynthesis process of streptomycin by the bacterium *Streptomyces griseus*, the dehydrogenation leading to the aldehyde formation is the last of 27 assembly steps, followed by the compound release from the bacteria for activation by StrA (Flatt et al., 2007). A putative gene product that would mediate this transition is unknown.

Comparison with the previous models of streptomycin-bound bacterial ribosomes further shows a discrepancy of the methyl group and the 6″ hydroxyl group conformations on the *N*-methyl-*L*-glucosamine moiety (Figure 1d). Our high-resolution structure provides the experimental evidence for the chemically more favorable conformation with the amino group at the 2″ position (protonated secondary amine) forming two hydrogen bonds/salt bridges with two backbone phosphates of rRNA, and the 6″ hydroxyl group can interact with one of the guanidino groups of the streptidine moiety (Figure 1d). Further, two water-coordinating magnesium ions are found (Figure 1c).

Overall, the streptomycin-bound structure on the SSU shows that potential modifications of medically relevant anti-bacterial compounds can be detected when bound to the human mitoribosome. In addition, specific water molecules and metal ions are revealed to be involved in the coordination. Since streptomycin is a drug that is used for the treatment of tuberculosis, the structural studies can be informative for designing less-toxic drugs.

## Materials and methods

### Sample preparation for cryo-EM

Flp-In T-Rex human embryonic kidney 293T (HEK293T) cell line (Invitrogen) was cultured as described previously (Khawaja et al., 2020), Penicillin-Streptomycin (Thermo Fisher) was supplemented to the media with the final concentration of 100 μg/mL streptomycin, and mitochondria were prepared as described previously (Aibara et al., 2018). The purified mitochondria were lysed by incubating at 4 °C for 20 min in the lysis buffer (25 mM HEPES-KOH pH 7.5, 5.0 mM Mg(OAc)_2_, 100 mM KCl, 2% (v/v) Triton X-100, 0.2 mM DTT, 1X cOmplete EDTA-free protease inhibitor cocktail (Roche), 40 U/μl RNase inhibitor (Invitrogen). Lysate was centrifuged at 5,000 g for 5 minutes at 4 °C and the supernatant was added to ANTI-FLAG M2 Affinity Gel (Sigma-Aldrich), equilibrated with the wash buffer (25 mM HEPES-KOH pH 7.5, 5.0 mM Mg(OAc)_2_, 100 mM KCl, 0.05% β-DDM). After 3 h incubation at 4 °C, the gel was washed with the wash buffer and the IF3-bound complexes were eluted by additional incubation of 2 h with the PreScission protease (GE Healthcare) (2 U/μl).

### Data collection and processing

3 μL of ~120 nM mitoribosome was applied onto a glow-discharged (20 mA for 30 sec) holey-carbon grid (Quantifoil R2/2, copper, mesh 300) coated with continuous carbon (of ~3 nm thickness) and incubated for 30 sec in a controlled environment of 100% humidity and 4 °C. The grids were blotted for 3 sec, followed by plunge-freezing in liquid ethane, using a Vitrobot MKIV (FEI/Thermofischer). Datasets were collected on Titan Krios (FEI/Thermofischer) transmission electron microscope operated at 300 keV, using C2 aperture of 70 μm and a slit width of 20 eV on a GIF quantum energy filter (Gatan). A K2 Summit detector (Gatan) was used at a pixel size of 0.83 Å (magnification of 165,000x) with a dose of 29-32 electrons/Å^2^ fractionated over 20 frames. A defocus range of −0.5 to −3.6 μm was used. More detailed parameters are listed in Table S1.

For the processing, movie frames were aligned and averaged by global and local motion corrections by RELION 3.0 (Zivanov et al., 2018). CTF parameters were estimated by Gctf (Zhang et al., 2016). Particles were picked by Gautomatch (http://www.mrc-lmb.cam.ac.uk/kzhang). The picked particles were subjected to 2D classification to discard contaminants as well as the LSU and monosome particles. The remaining particles underwent 3D auto-refinement with RELION 3.0 using EMD-10021 as a 3D reference, followed by 3D classification with local angular search with a solvent mask to remove poorly aligned particles. Well-resolved classes were pooled and subjected to 3D refinement and CTF refinement (beam-tilt, per-particle defocus, per-micrograph astigmatism) by RELION 3.1 (Zivanov et al., 2020), followed by Bayesian polishing. Particles were then separated into multi-optics groups based on acquisition areas and date of data collection. Second round of CTF refinement (beam-tilt, trefoil and fourth-order aberrations, magnification anisotropy, per-particle defocus, per-micrograph astigmatism) was performed, followed by 3D auto-refinement. To improve the local resolution, local-masked 3D auto-refinements were performed (Figure 1—figure supplement 1).

Reported resolutions are based on applying the 0.143 criterion on the FSC between reconstructed half-maps. Finally, the maps were subjected to *B*-factor sharpening and local-resolution filtering by RELION 3.1, superposed to the overall map and combined for model refinement.

### Model building and refinement

For the SSU bound with streptomycin, the starting model was PDB ID: 6RW4. Manual revision was done using *Coot* 0.8 (Emsley et al., 2010). The streptidine (inositol with two hydroxyl groups substituted by guanidino groups) and the *N*-methyl-*L*-glucosamine parts agreed with the density, while the aldehyde group of streptose part disagrees. Therefore, the hydrated aldehyde was placed based on the reported hydration information (Blundell et al., 2013). The water molecules were automatically picked by *Coot*, followed by manual revision. Geometrical restraints of modified residues and ligands were calculated by Grade Web Server (http://grade.globalphasing.org) or obtained from CCP4 library (Lebedev et al., 2012). Hydrogens were added to the models except for water molecules by REFMAC5 (Murshudov et al., 2011) using the prepared geometrical restraint files. The model was then refined against the composite map using Phenix.real_space_refinement v1.18 (Liebschner et al., 2019) with global energy minimization with reference restraints (only for non-modified protein residues, using the input model as the reference, sigma 5) and rotamer restraints, without Ramachandran restraints. Validation was done by MolProbity (Williams et al., 2018). The statistics are listed in Table 1.

## Accession codes

The cryo-EM density maps and atomic coordinates have been deposited in the Electron Microscopy Data Bank (EMDB) and Protein Data Bank (PDB) under accession codes EMD-13170 and 7P2E.

## Acknowledgements

Cryo-EM data were collected at the cryo-EM Swedish National Facility (funded by KAW, EPS, and Kempe foundations) at SciLifeLab. This work was supported by the Swedish Foundation for Strategic Research (FFL15:0325), Ragnar Söderberg Foundation (M44/16), European Research Council (ERC-2018-StG-805230), Knut and Alice Wallenberg Foundation (2018.0080).

---

**Figure 1—figure supplement 1.**
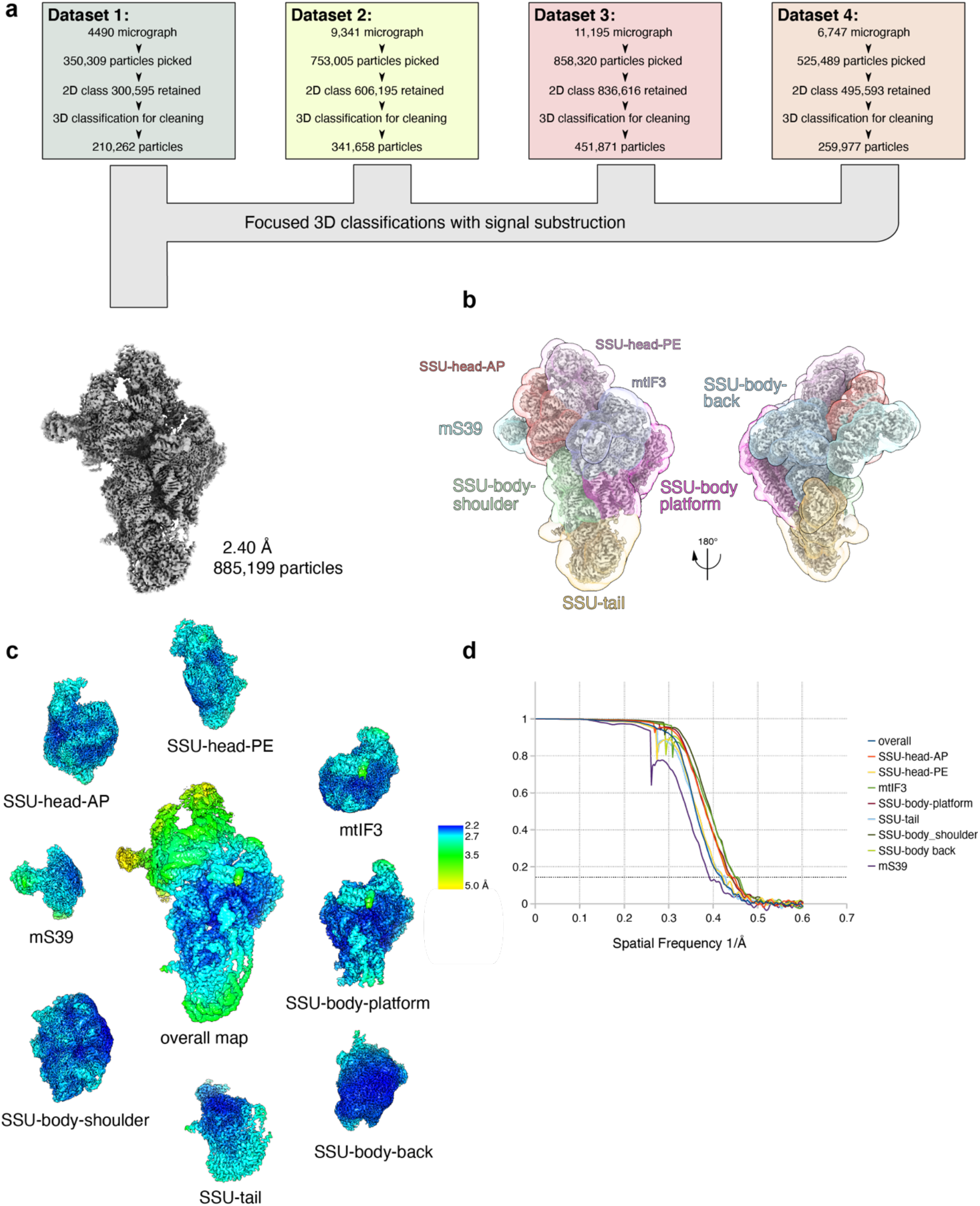
Cryo-EM data collection and processing. (a) Cryo-EM data processing overview. (b) Binary masks used for local-masked refinements. (c) Density maps from overall refinements, filtered by their nominal resolution without sharpening, colored by local resolution. (d) Fourier shell correlation (FSC) curves of half maps from overall refinement and the local-masked refinements.

